# Sport expertise and motor imagery abilities shape sensorimotor rhythm modulations during visualisation tasks: Implications for neurofeedback-based cognitive training in athletes

**DOI:** 10.64898/2026.08.26.747187

**Authors:** Margaux Izac, Emeline Pierrieau, Eléa Rossignol, Natalia Grechukhin, Elina Coudroy, Léa Pillette, Bernard N’Kaoua, Camille Jeunet-Kelway

**Author notes:** Author to whom any correspondence should be addressed. These authors contributed equally (Margaux Izac, Emeline Pierrieau). **Declaration of interest:** The authors declare that they have no known competing financial interests or personal relationships that could have appeared to influence the work reported in this paper.

## Abstract

Kinaesthetic motor imagery (kMI) is widely used in sport to enhance motor performance by engaging cortical sensorimotor networks. Neurofeedback may further support kMI, but the optimal neural target to reinforce remains unclear. Maximal sensorimotor event-related desynchronisation (SMR-ERD) represents a relevant target as it may index sensorimotor cortex engagement, yet sport expertise has been associated with reduced SMR-ERD, potentially reflecting neural efficiency. The optimal neurofeedback target may therefore depend on sport expertise, movement expertise, and individual kMI ability. This study examined how these factors influence sensorimotor activity during kMI. We compared 17 basketball players (Experts) and 16 individuals without formal basketball training (Novices). kMI ability and frequency of use were assessed using questionnaires, while SMR-ERD was quantified using electroencephalography (EEG) during kMI. Participants imagined either a basketball-specific movement (Free throw), for which only Experts had extensive experience, or a generic movement (Box lifting), familiar to both groups. Experts reported greater kMI ability and more frequent kMI use than Novices. Only Experts exhibited significant and sustained SMR-ERD during kMI. Moreover, SMR-ERD was stronger in Experts than Novices specifically during Free throw kMI, corresponding to their movement of expertise. Nonetheless, within the Expert group, higher kMI ability was associated with reduced SMR-ERD. These findings suggest that sport expertise initially enhances voluntary recruitment of sensorimotor networks during kMI, whereas greater kMI ability may subsequently promote neural efficiency, resulting in reduced overall sensorimotor cortical activation. These results highlight the need to tailor kMI-based neurofeedback training to users’ sport expertise and kMI ability levels.

**Highlights:** · Current motor imagery-based neurofeedback designs may not be tailored to athletes.

· Athletes show greater recruitment of sensorimotor cortex during motor imagery.

· High motor imagery ability in athletes may be associated with neural efficiency.

· Neurofeedback should be adapted to sport expertise and motor imagery ability.

## 1. Introduction

Motor imagery (MI) can be defined as a “dynamic state during which one simulates an action mentally without any body movement” (Guillot et al., 2008). Because it requires no physical effort, MI represents an effective complement to physical training: it helps prevent excessive training loads, maintain training during injury periods, and supports athletes’ physical and mental well-being (Guillot & Collet, 2008). Previous research has demonstrated the positive impact of MI, and particularly of kinaesthetic MI (kMI), which consists in recalling proprioceptive sensations, including tactile sensations, body temperature, muscle contraction or relaxation, experienced during physical action execution, on motor skills, including improvements in precision (Deng et al., 2024), flexibility (Battaglia et al., 2014), strength (Piveteau et al., 2023), and speed (Gentili & Papaxanthis, 2015). These benefits rely on the fact that physical execution and kMI engage overlapping motor-related brain regions (Dechent et al., 2004; Filgueiras et al., 2018; Guillot et al., 2008; Hanakawa et al., 2008; Hardwick et al., 2018; Henschke & Pakan, 2023; Hétu et al., 2013; Raffin et al., 2012). However, despite its demonstrated benefits, a major limitation of kMI training is the absence of indicators of performance. Indeed, unlike physical training, during kMI movement-related brain activity cannot be adjusted on the basis of sensory information, as no overt motor output is produced. This lack of feedback may hinder motivation and learning (Abbas & North, 2018; Ericsson et al., 1993; Sigrist et al., 2013).

Electroencephalography (EEG) offers a means to detect the engagement of sensorimotor cortical regions during kMI. Indeed, movement execution and, to a lesser extent, kMI, are accompanied by a transient decrease in the power of sensorimotor rhythms (SMR), particularly in the mu (8-12 Hz) and low beta (12-20 Hz) frequency bands, over the sensorimotor cortex. This phenomenon, referred to as SMR event-related desynchronisation (SMR-ERD) (Nojima et al., 2022; Soekadar et al., 2015), is thought to reflect the activation of underlying cortical neurons (Formaggio et al., 2010; Zich et al., 2015). SMR-ERD magnitude can therefore serve as an index of the recruitment of sensorimotor cortical areas during kMI (Neuper et al., 2006). Neurofeedback (NF) training can be used to exploit these known features by translating SMR-ERD magnitude into an interpretable sensory cue (e.g., auditory or visual feedback), providing users with near-real-time feedback on their ability to self-regulate target brain activities, and thereby adjust their strategy to maximise the engagement of their sensorimotor cortical areas during kMI (Miller et al., 2010). An increasing number of studies support that this closed-loop training enhances athletes’ ability to self-regulate targeted brain activities, and thereby sport performance (Gong et al., 2020; Gruzelier et al., 2014; Wu, Chueh, et al., 2024; Wu, Tu, et al., 2024), making it a promising tool to augment mental preparation in sports (Debarnot et al., 2014; Gong et al., 2021; Mirifar et al., 2017; Xiang et al., 2018; Yu et al., 2025).

Across domains, most MI-based NF (MI-NF) protocols provide feedback that rewards maximal SMR-ERD (Horowitz et al., 2021). While this approach may benefit individuals who have difficulty engaging sensorimotor regions, its suitability for athletes can be questioned. Although evidence on the effects of sport expertise on SMR-ERD remains limited, several studies suggest that athletes generate more consistent SMR-ERD during MI than novices (Sugino & Ushiyama, 2021; Wolf et al., 2014), and show reduced SMR-ERD during movement execution (Del Percio et al., 2010) and action judgment tasks (Babiloni et al., 2010; Del Percio et al., 2009). This attenuation of SMR-ERD may reflect lower sensorimotor cortical activation when athletes perform movements related to their sport expertise (Chang et al., 2011; Kim et al., 2014; Kraeutner et al., 2018; Li & Smith, 2021; Wriessnegger et al., 2022), consistent with neural efficiency mechanisms (Haier et al., 1988; Li & Smith, 2021) resulting from extensive movement practice. However, it remains unclear whether these neural efficiency mechanisms also apply during kMI, resulting in a parallel attenuation of SMR-ERD when athletes imagine sport-related movements. Importantly, reduced SMR-ERD during kMI could also reflect superior kMI ability, as athletes typically engage in kMI more frequently than novices (Arvinen-Barrow et al., 2007; Gregg et al., 2011; Hall et al., 1990) and may therefore develop expertise in both the imagined movements and the imagery process itself. Finally, it remains unclear whether SMR-ERD attenuation observed in athletes is specific to movements performed in their sport or also extend to biomechanically similar, sport-unrelated movements, given that movement expertise is known to transfer across related motor skills. Clarifying these relationships is essential for developing MI-NF protocols tailored to optimize athletes’ motor performance.

Accordingly, the present study investigated how SMR-ERD during kMI, as measured in typical MI-NF protocols, is influenced by sport expertise (athletes vs. novices), movement expertise (sport-related vs. sport-unrelated movements), and individual differences in kMI ability and frequency of use.

To this end, we recorded EEG during kMI in a group of 17 basketball players (Experts) and 16 non-basketball players with no regular sport practice (Novices). We included both an expertise-specific kMI task (Free throw) and a generic kMI task (Box lifting). We also assessed individual kMI frequency of use and kMI ability using self-report questionnaires (Hall et al., 1990; Robin et al., 2020) prior to the experimental task. We hypothesised that Experts would exhibit greater kMI ability and more frequent use, as well as a superior capacity to voluntarily generate SMR-ERD during kMI compared to Novices. In line with the neural efficiency framework, we further expected that the association between SMR-ERD amplitude and kMI ability would differ between groups, such that Experts with higher kMI ability would show more efficient sensorimotor activation, reflected in reduced SMR-ERD during kMI.

## 2. Materials and Methods

### 2.1. Participants

Thirty-three healthy participants were recruited and assigned to two groups: an expert group (Experts; n = 17) of basketball players (age = 20.6 ± 2.4 years; 9 women, 8 men) and a novice group (Novices, n = 16) of non-basketball players (age = 22.7 ± 3.8 years; 8 women, 8 men). Basketball players sport practice was quantified using three indices. Weekly practice frequency was rated on a 4-point scale 1 = “*from time to time*”, 2 = “*once per week*”, 3 = “*several times per week*”, and 4 = “*every day*” (3.3 ± 0.9). Years of basketball experience averaged 11.6 ± 3.9 years. was evaluated on a 4-point scale: 1 = Departmental level, 2 = Regional level, 3 = National level, and 4 = Feminine/Professional league (2.1 ± 0.8). Detailed scores are provided in Supplementary Materials. Novices reported no prior formal basketball training and no substantial expertise in other sports, musical activities, or video gaming that could involve advanced kMI abilities. All participants were naïve to NF interventions. The Edinburgh Handedness Inventory (Oldfield, 1971), revealed that 28 participants were right-handed (mean laterality score = 88.68%) and 4 were left-handed (mean laterality score = -77.38%).

### 2.2. Experimental design

Participants were seated in front of a 27-inch computer screen and completed a series of questionnaires. They were then presented with a general explanatory video detailing the experimental protocol. Participants were informed that EEG recordings are highly sensitive to muscular artefacts and were therefore instructed to remain as still as possible throughout the trials. They were also familiarised with the kMI modality, which consists in recalling the sensations perceived (e.g., tactile sensations, body temperature, muscle contraction or relaxation) during the execution of a movement, as it predominantly recruits parietal and motor-related regions (Guillot et al., 2009; Kwon et al., 2023; Solodkin et al., 2004).

The experiment consisted of two blocks, each corresponding to one imagined task (i.e., Box lifting or Free throw), for which the order was counterbalanced across participants. Each block began with the presentation of a video depicting a 3D stick-avatar executing an action. The actions were either a basketball free throw (Free throw) or a box lifting action (Box lifting). Both tasks involved whole-body movements. The Free throw was selected because it is a widely practiced basketball skill that can be consistently executed within 10 seconds. The Box lifting consisted of moving a cardboard box, between a knee height shelf and a higher shelf located sufficiently high to require standing on their toes. This task was designed to be biomechanically similar to a free throw while familiar to all participants. During a familiarisation phase, participants were provided with a basketball or a cardboard box and instructed to physically perform the task shown in the video. They were instructed to progressively reduce movement amplitude until adopting a seated position and engaging exclusively in kMI. Once participants were ready, they completed the first of 4 runs, each comprising 10 kMI trials. A run lasted approximately 3min. Each single run began with a 30s resting state period, followed by 10 trials for which EEG activity was recorded. Each trial comprised a 2s baseline, a 10s kMI period, and a variable rest period of 1-3s preceding the next trial’s baseline (Figure 1). Start and stop cues were provided via auditory beeps and through the display of a white fixation cross on a black background.

**Figure 1.**
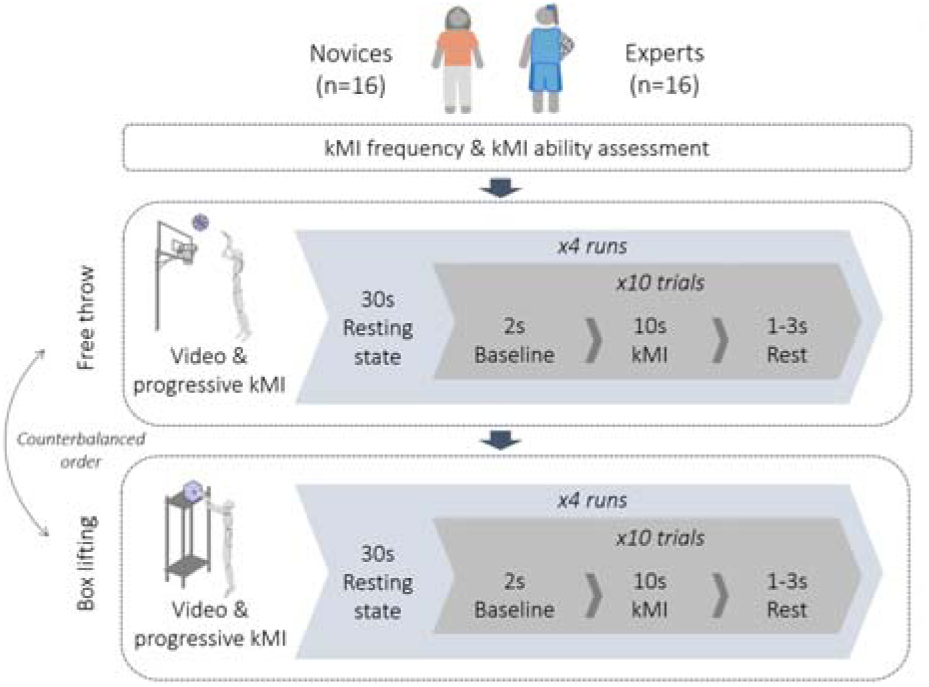
Experimental Paradigm. The session began with questionnaire data collection. It was then divided in two blocks, one for each task. A block started with a 3D stick avatar video showing the task to imagine (Free throw or Box lifting). Participants were asked to execute the task and progressively engage in kMI. A run then started with a 30s resting state immediately followed by 10 trials. One trial consisted of a 2s baseline, a 10s kMI period and 1 to 3s resting phase. A block was complete when 4 runs were recorded.

Participants were instructed to fixate the centre of the screen throughout the experiment. During kMI periods, participants were instructed to imagine executing the task visualised in the video once. If the mental representation was completed before the cross on screen disappeared, participants were asked to repeat the imagined action a second time within the same trial. In subsequent trials, participants were encouraged to better regulate the speed of their imagery.

Short rest periods were provided between runs and blocks and were terminated when participants felt ready to proceed.

### 2.3. Questionnaire data

A modified version of the Imagery Use Questionnaire (IUQ) (Hall et al., 1990) was first administered to assess kMI frequency of use, rated on a 7-point Likert scale ranging from *Never* to *Always*. Participants then completed the French version of the Motor Imagery Questionnaire-Third version (MIQ-3f) (Robin et al., 2020) to evaluate MI ability, rated on a 7-point Likert scale ranging from *Very difficult* to *Very easy*. For both questionnaires, given that participants were instructed to use kinaesthetic strategies during MI tasks, only items assessing kMI were retained, namely kMI ability and kMI frequency of use.

### 2.4. EEG recordings and pre-processing

EEG was recorded using an ANT Neuro eego™sports gel-based headset with a 32-channel montage positioned according to the international 10-20 system (FP1, FPz, FP2, F7, F3, Fz, F4, F8, FC5, FC1, FC2, FC6, A1, T7, C3, Cz, C4, T8, A2, CP5, CP1, CP2, CP6, P7, P3, Pz, P4, P8, POz, O1, Oz, O2), and amplified using eego™sports or eego™rt amplifiers. Signals were recorded with an online reference at CPz and a ground electrode at AFz, and electrode impedances were maintained below 5 kΩ. EEG signals were recorded via OpenVibe (Renard et al., 2010) and pre-processed using EEGLAB (Delorme & Makeig, 2004) and Fieldtrip (Oostenveld et al., 2010) in Matlab.

Data were first downsampled to 250Hz to reduce computational load while maintaining adequate temporal resolution for the frequency range of interest. Data were then band-passed filtered between 1 and 45Hz using a zero-phase (two-pass) finite impulse response (FIR) filter to remove slow drifts and high-frequency noise. Noisy channels were automatically detected using the Clean Rawdata algorithm from EEGLAB. Channels were rejected if they were flat for more than 5s, their line-noise level exceeded a robust z-score of 4 relative to other channels, or showed a correlation with neighbouring channels that was inferior to 0.8. Rejected channels were removed and subsequently interpolated based on the activity from their neighbouring electrodes, after which the EEG data were re-referenced to the common average reference in order to further minimise noise. Continuous EEG was then segmented into 12.5s epochs, starting 2.5s before the display of the cross on the screen (baseline period) and ending 0.5s after its disappearance (kMI period and beginning of rest). Finally, EEG data were transformed into current source density estimates using a spherical spline surface Laplacian as implemented in EEGLAG (function *pop_currentdensity*, spline method) to reduce volume conduction effects and enhance the spatial specificity of sensorimotor oscillatory activity.

Following pre-processing, Fieldtrip toolbox was used to apply time-frequency decomposition using Morlet wavelets (4-40 Hz with 1 Hz steps). Wavelet cycles were increased by 0.1 at each frequency, starting from a width of 3 to 6.6 to ensure a balance between sufficient temporal resolution at lower frequencies and frequency resolution at higher frequencies. Baseline correction was applied using the average power during the 2s pre-cue period as baseline and a dB normalisation to facilitate comparisons across trials and participants. Data were then downsampled to a total of 200 time points in order to avoid memory issues while ensuring adequate temporal resolution for graphical displays.

### 2.5. Analysis

Due to a technical issue in EEG recordings, one participant was excluded from all analyses. As a result, each group comprised 16 participants. Two Expert participants had 2 and 10 trials missing, respectively; given the low proportion of missing data (2.5% and 12.5%), these participants were retained in the analyses.

Statistical analyses were computed using Jamovi v.2.6.19 (https://www.jamovi.org), which implements R statistical language (https://www.R-project.org/).

For all statistical analyses, normality was first assessed using Shapiro*-*Wilk test, and homogeneity of variances was evaluated using Levene’s test. As the data met the assumptions of normality and homoscedasticity (p > 0.05 at Shapiro-Wilk and Levene’s tests), independent-samples t-tests were conducted for between-group comparisons (Experts vs. Novices), and paired-samples t-tests were used for within-group comparisons (Free throw vs. Box lifting). The significance of SMR-ERD was examined using one-sample t-tests against zero separately for Experts and Novices. Effect sizes for all t-tests were calculated using Cohen’s d, with values of 0.2, 0.5, and 0.8 interpreted as small, medium, and large effects, respectively (Cohen, 1988).

The effects of between- and within-group factors on SMR-ERD were analysed using a repeated-measures ANOVA comparing mean power change across the entire kMI period, with Group (Novices vs. Experts) as a between-subject factor and kMI task (Box lifting vs. Free throw) as a within-subject factor. Effect sizes for ANOVA results were reported as partial eta-squared (η²p). Post hoc analyses were conducted using paired- or independent-samples t-tests, as appropriate. When multiple comparisons were performed, p-values were adjusted using the False Discovery Rate (FDR) correction according to the Benjamini-Hochberg procedure (Benjamini & Hochberg, 1995).

Finally, the relationships between SMR-ERD and kMI ability and kMI frequency of use were assessed respectively in both Experts and Novices using Spearman’s rank-order correlation.

## 3. Results

### 3.1. Self-reported kMI abilities and frequency of use in Experts vs. Novices

Mean score obtained for kMI frequency of use and kMI ability were compared between Novices and Experts. A significant group difference was found for kMI frequency of use (t(30) = -2.95, p = 0.006, d = 1.04), with Experts reporting more frequent use of kMI than Novices. Consistent with these findings, results also indicated higher kMI ability scores in Experts compared to Novices (t(30) = -2.09, p = 0.045, d = 0.74). Those results are illustrated in Figure 2.

**Figure 2.**
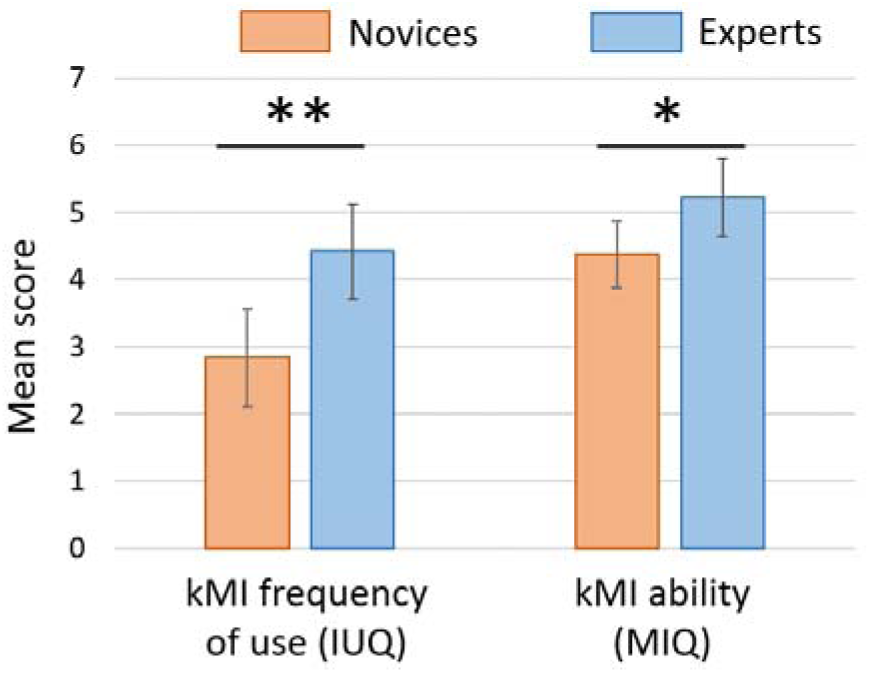
Mean kMI frequency (left) and kMI ability (right) scores in Novices (orange) and Experts (blue). Error bars indicate 95% confidence intervals. *p < 0.05, **p < 0.01

### 3.2. SMR-ERD magnitude during kMI tasks in Experts vs. Novices

The next step of the analyses examined whether Novices’ and Experts’ SMR-ERD differed during kMI. SMR-ERD was quantified by averaging power relative to the 2s pre cue baseline, within the 8-15 Hz frequency band, as the strongest power changes from baseline over central electrodes (C3, Cz, C4) were observed in this range for both groups (Figure 3A and 3B).

**Figure 3.**
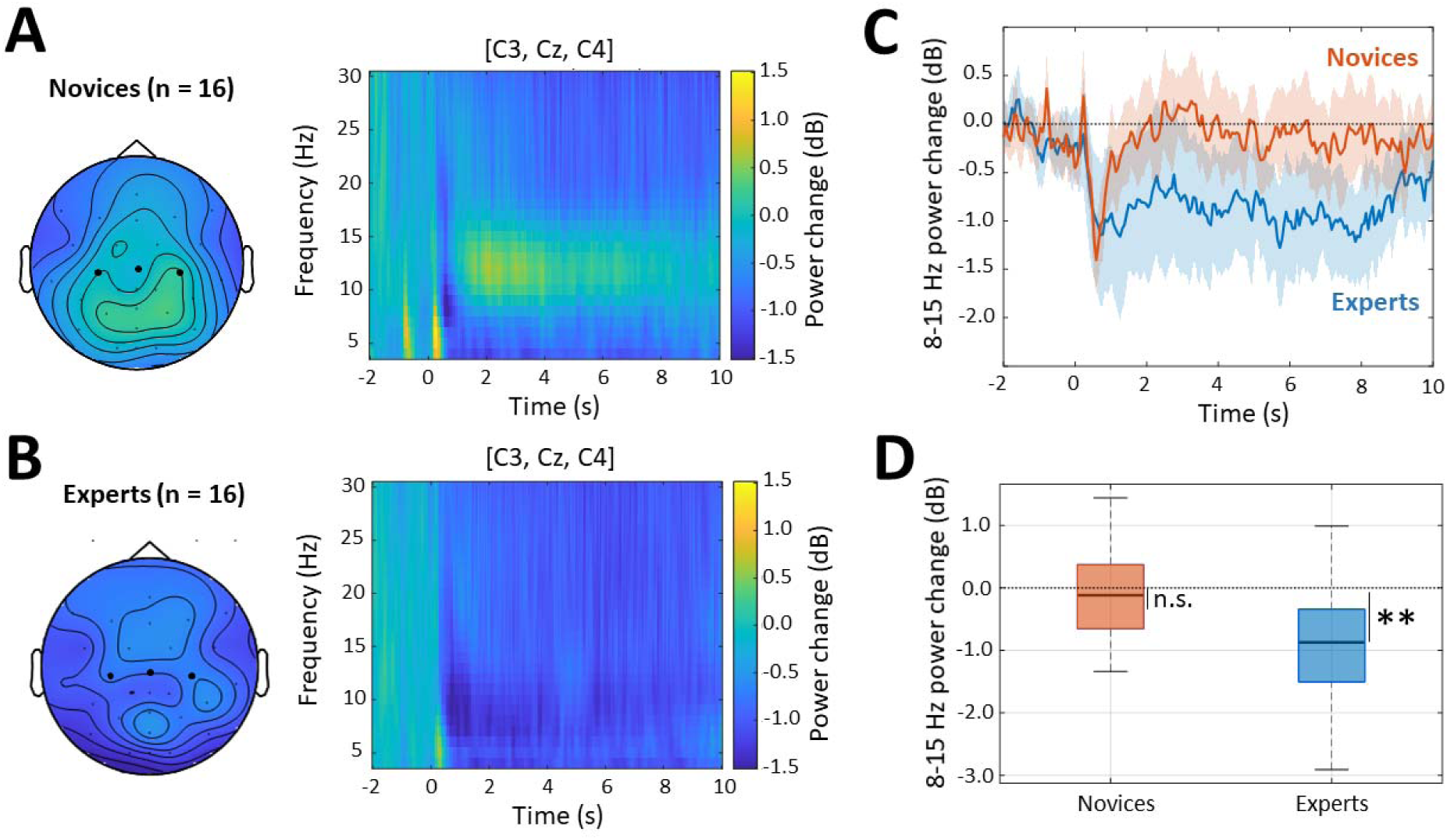
SMR-ERD in Novices and Experts. **A**, **B**. Topographical maps represent the average 8-15 Hz power change (dB) during kMI across electrodes, with C3, Cz, and C4 highlighted in bold, and time-frequency maps represent power change over central electrodes (C3, Cz, C4) specifically, in Novices (A) and in Experts (B). Blue and yellow colours respectively indicate a decrease and increase in power relative to baseline (pre-kMI) activity. **C**. 8-15 Hz power change over central electrodes (C3, Cz, C4) during kMI (t0 = kMI cue onset) in Novices (orange) and in Experts (blue). Shaded areas represent 95% confidence intervals. **D**. Mean 8-15 Hz power change over central electrodes (C3, Cz, C4) during the entire kMI duration in Novices (orange) and in Experts (blue). **p < 0.01, n.s. = not significant.

Novices exhibited a transient SMR-ERD (i.e., decrease in 8-15 Hz power over central electrodes) during the first second following kMI cue onset, after which activity returned to baseline (pre-cue) levels for the remainder of the kMI period (Figure 3C, orange line). In contrast, Experts demonstrated a sustained SMR-ERD that persisted throughout the 10s kMI period (Figure 3C, blue line). These differences were confirmed by statistical analyses (Figure 3D): when averaged across the entire kMI duration, mean power was significantly below zero in Experts (t(15) = -3.79, p = 0.002, d = -0.95), indicating significant SMR-ERD. In contrast, mean power did not significantly differ from zero in Novices (t(15) = -0.43, p = 0.671, d = -0.11).

### 3.3. SMR-ERD magnitude during kMI of generic vs. sport expertise-specific tasks

To examine whether SMR-ERD during kMI was further modulated by movement expertise, we compared SMR-ERD between Novices and Experts while they imagined either a movement for which neither group had specific expertise (Box lifting) or a movement for which only Experts had specific expertise (Free throw).

When examined across the time course of kMI trials, the absolute difference in SMR-ERD between Experts and Novices appeared more pronounced for Free throw than for Box lifting (dark vs. light coloured lines in Figure 4A). This observation was confirmed by a repeated-measures ANOVA which revealed a significant main effect of Group (F(1,30) = 10.7, p = 0.003, η^2^p = 0.26), no significant main effect of kMI task (F(1,30) = 0.1, p = 0.730, η^2^p = 0.00), and a significant Group × kMI task interaction (F(1,30) = 5.6, p = 0.025, η^2^p = 0.16). Following FDR correction for multiple comparisons, only the post hoc comparison between Novices and Experts in the Free throw condition remained significant, with Experts showing significantly lower power (i.e., stronger SMR-ERD) than Novices (t(30) = -3.65, p = 0.004, d = -1.29) (Figure 4B). Thus, when compared to Novices, Experts specifically showed greater SMR-ERD during kMI of movements directly related to their sport expertise (i.e., Free throw).

**Figure 4.**
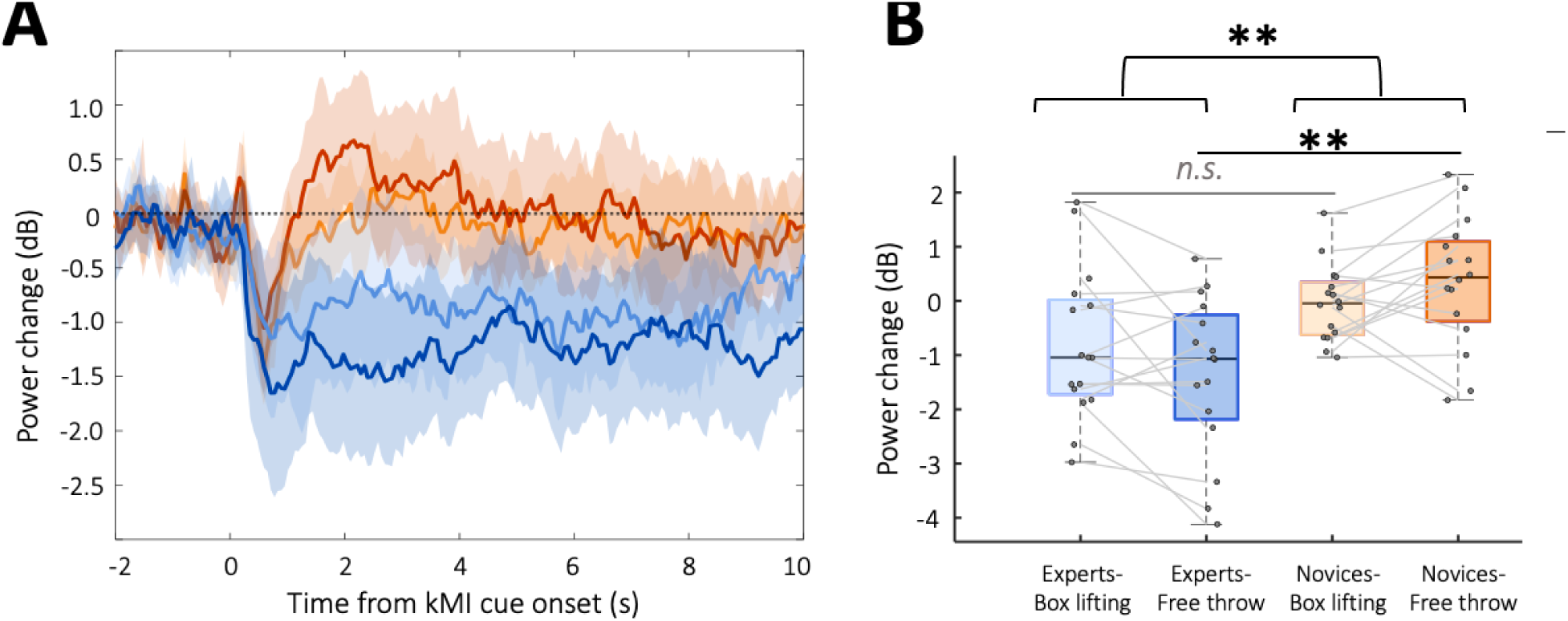
SMR-ERD according to Group (Novices vs. Experts) and kMI task (Box lifting vs. Free throw). **A**. Time course of power change relative to kMI cue onset. Each condition is represented as a coloured line (light orange = Novices / Box lifting, dark orange = Novices / Free throw, light blue = Experts / Box lifting, dark blue = Experts / Free throw). Shaded areas indicate 95% confidence intervals. **B**. Mean power change across the entire kMI period. Connected dots illustrate individual data pertaining to the same participants. **p < 0.01, n.s. = not significant

### 3.4. Relationship between kMI ability, frequency of use, and SMR-ERD in Experts and Novices

The association between SMR-ERD and self-reported kMI ability/frequency of use scores was examined by using correlation analyses. Results showed a significant positive correlation between SMR power change and kMI ability scores in Experts (Spearman’s ρ = 0.50, p = 0.047), whereas the correlation was not statistically significant in Novices (Spearman’s ρ = -0.36, p = 0.165) (Figure 5). The same correlation analyses were carried for SMR-ERD and kMI frequency of use but yielded no significant results for both groups. Thus, within the Experts group only, individuals reporting the highest kMI ability were those showing the smallest SMR-ERD during kMI.

**Figure 5.**
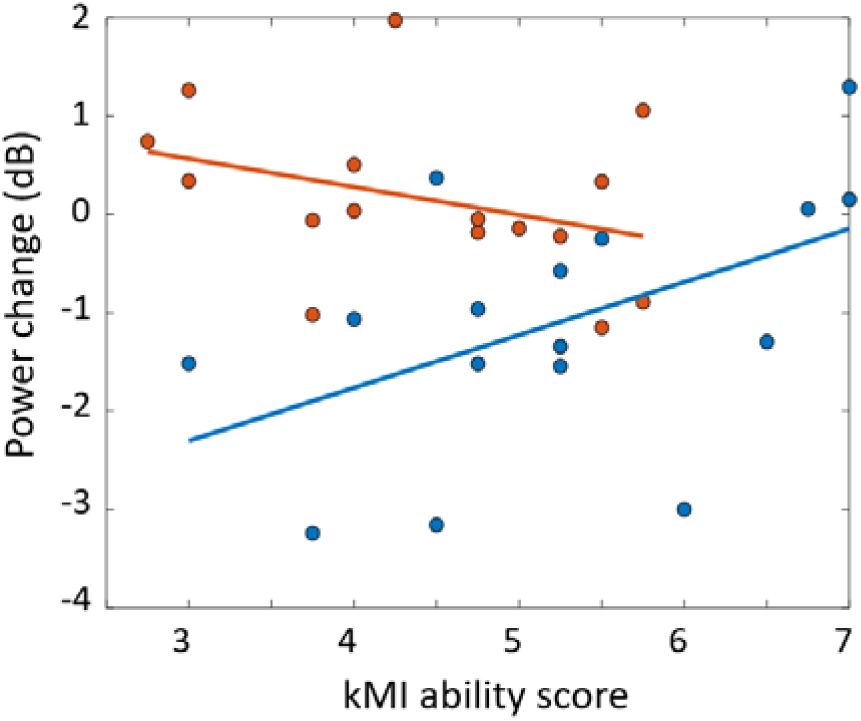
Correlation between kMI ability score and power change. Orange and blue dots indicate individual data from Novices and Experts, respectively. Regression lines indicate the direct of the linear relationship between the two variables in Experts (blue line) and in Novices (orange line).

## 4. Discussion

The objective of the present study was to investigate the modulatory roles of sport expertise, movement expertise, and kMI ability/frequency of use on SMR-ERD during kMI in order to guide the design of relevant and efficient MI-NF cognitive training procedures for athletes. More precisely, to characterise the potential neural efficiency phenomenon, we compared SMR-ERD magnitude in 17 basketball players (Experts) and 16 non-basketball players (Novices) while they performed kMI of a task mastered only by Experts (Free throw) or a task requiring no particular skill (Box lifting). Additionally, the participants’ kMI ability and frequency of use were assessed through validated questionnaires.

Experts reported higher kMI ability and more frequent use of kMI than Novices. In addition, only Experts demonstrated significant SMR-ERD during kMI. Including movement expertise as a factor in the analyses revealed that this difference in SMR-ERD was significant only during Free throw kMI, indicating that Experts engaged sensorimotor cortical regions more strongly than Novices specifically when imagining tasks in which they had developed movement expertise. Furthermore, a significant correlation between kMI ability scores and SMR power change during kMI was observed only within the Experts group, with higher kMI ability being associated with smaller SMR-ERD.

The finding of Experts, and not Novices, being able to voluntarily engage their sensorimotor cortex during kMI, as reflected in significant SMR-ERD, aligns with previous studies that showed greater neural recruitment and excitability within task-relevant regions with sport expertise (Pfurtscheller, 2000). The better ability of Experts to produce a reliable and sustained SMR-ERD during kMI may also be partly explained by differences in kMI ability and frequency of use. In the current sample, Experts reported higher kMI ability and more frequent engagement in kMI practice, consistent with previous findings (Bek et al., 2025; Gregg et al., 2011; Hall et al., 1990; Mao et al., 2022; Williams et al., 2015). MI ability has been identified as a key determinant of sensorimotor engagement during MI. For instance, an fMRI study reported that individuals with high MI ability manifested greater activation in the parietal and ventrolateral premotor region during MI compared to low-ability imagers (Guillot et al., 2008). Similarly, research using MI-NF protocols has demonstrated that successful classification of left-versus right-hand MI is strongly dependent on MI ability, with skilled imagers producing more robust and discriminable SMR-ERD patterns than poor imagers, whose SMR-ERD tends to be weaker and less consistent (Marchesotti et al., 2016; Vuckovic & Osuagwu, 2013). Furthermore, in the present experiment, the Free throw represented an action in which only Experts had extensive practice. Consistently, the difference in the ability to induce SMR-ERD during kMI in Experts and Novices was specifically significant in this expertise-specific condition, and not in the Box lifting task, which required no particular skill. This finding supports the hypothesis that well-established motor representations, acquired through extensive physical training, facilitate the recruitment of sensorimotor networks during MI and thus the generation of SMR-ERD (Olsson & Nyberg, 2010). Accordingly, the distinct capacity in generating SMR-ERD between Experts and Novices likely comes from differences in sport expertise and kMI ability, both contributing to the ability to actively engage sensorimotor regions during kMI.

When examining the relationship between SMR-ERD and kMI ability, correlation analyses revealed that, within the Experts group only, higher kMI ability was associated with smaller SMR-ERD, suggesting that better imagers produced less sensorimotor cortical activation during kMI. This finding is in line with the involvement of neural efficiency mechanisms during kMI in Experts. According to the neural efficiency framework, individuals with higher levels of expertise tend to recruit task-relevant motor networks more selectively and economically, particularly once a high degree of proficiency has been achieved (Haier et al., 1988; Li & Smith, 2021). Several EEG studies have demonstrated reduced SMR-ERD in expert athletes compared to novices across various conditions (Babiloni et al., 2010; Del Percio et al., 2009). For instance, Del Percio et al. (2010) reported that wrist extensions elicited lower 8-10 Hz ERD in primary motor and premotor regions in experts compared with novices, during both movement preparation and execution. Similarly, movement-related cortical potentials were reduced in expert athletes relative to novices during finger flexions (Di Russo et al., 2005) or observation of sport-specific actions and key pressing (Del Percio et al., 2008). Smaller but consistent SMR-ERD may index more efficient recruitment of sensorimotor regions. Our results extend previous work by suggesting that neural efficiency may emerge through a two-step process. First, sport expertise appears to enhance the ability to recruit sensorimotor networks during kMI, as reflected by the capacity of Experts, but not Novices, to generate significant SMR-ERD. This reliable recruitment may constitute a prerequisite for the emergence of neural efficiency. Once established, sensorimotor activation may become optimised, leading to more energetically efficient engagement during kMI. This interpretation is consistent with Debarnot et al. (2014), who proposed that motor expertise and repeated imagery practice contribute to the progressive refinement of internal motor representations.. Thus, the combination of sport and kMI expertise may provide athletes with richer and more refined sensorimotor representations, enabling more precise engagement of relevant neural circuits during kMI.

By highlighting a potential two-step process, our findings may help explain why some studies report increased cortical activations and others reductions with sport expertise during MI. Variability in MI ability within expert samples could lead to distinct patterns of sensorimotor engagement: studies including a higher proportion of high kMI ability experts may report weaker activations, whereas those with more low-ability imagers may report stronger activations. However, this interpretation remains difficult to confirm, as MI ability is not systematically assessed in neuroimaging studies comparing individuals of varying sport expertise level during MI (Kim et al., 2014; Wolf et al., 2014; Wriessnegger et al., 2022).

Importantly, these findings shed light on MI-NF applications aimed at enhancing motor performance in athletes. While MI-NF has often yielded positive results in clinical contexts to increase SMR-ERD and enhance motor recovery (Cha et al., 2025; Pichiorri et al., 2015), the application of MI-NF in sport settings has yielded mixed findings (Xiang et al., 2018; Yu et al., 2025). The present results suggest that this inconsistency may stem from a lack of adaptation of the rewarded brain activity pattern to individuals’ sport expertise and kMI ability. More precisely, our findings suggest that the classical MI-NF paradigms, aiming to train individuals to reliably engage sensorimotor networks during kMI through reinforcement of SMR-ERD, may not be optimal in athletes, since most of them may already master this capacity. Based on our findings, some Experts may even benefit from MI-NF protocols encouraging reduction of SMR-ERD amplitude during kMI, with the aim to optimise sensorimotor activation. Indeed, kMI ability scores were heterogenous within the Experts group and the negative correlation between SMR-ERD amplitude and kMI ability suggests that Experts with lower kMI ability may benefit from reduced SMR-ERD. However, the correlation found between SMR-ERD and kMI ability in this study was conducted on a limited sample size and of modest amplitude, suggesting that these results remain to be replicated and further deepen. For instance, future studies may benefit from including a broader spectrum of expertise levels, both in sport and kMI ability. This would enable to determine whether sport expertise and kMI ability are linearly related, or whether they represent partially independent processes, potentially giving rise to distinct athlete profiles with varying levels of kMI proficiency. Should a non-linear relationship between SMR-ERD magnitude and expertise be confirmed, MI-NF protocols will need to adapt the rewarded neural target to account for differences in sensorimotor cortical activation across individuals.

## 5. Conclusion

The present study indicates that sport expertise facilitates the recruitment of sensorimotor networks during kMI of skilled movements, as reflected by significant SMR-ERD in Experts and not in Novices. Higher kMI ability in Experts was associated with smaller SMR-ERD, consistent with the involvement of neural efficiency mechanisms according to kMI ability in Experts. These findings suggest that MI-NF interventions aiming to enhance motor performance in athletes should be adapted to the user’s level of sport expertise and kMI ability.

## Data availability statement

The data supporting the findings of this study are available from the corresponding author upon request.

## Conflict of interest

All authors declare having no competing/conflicts of interest.

## Acknowledgements

This work received financial support from the French government in the framework of the University of Bordeaux’s IdEx “Investments for the Future” program / GPR BRAIN2030 as well as the programme ANR-21-EXES-0004 - France 2030.

## Ethics statement

After being informed of the research aims, conditions and financial compensation, all participants gave their informed written consent. This research was approved by the French Protection of Persons’ Committee (national number 2022-A00626-37).

## Notes

### Competing Interest Statement

The authors have declared no competing interest.

