## Supplementary Material for "Sport expertise and motor imagery abilities shape sensorimotor rhythm modulations during visualisation tasks: Implications for neurofeedback-based cognitive training in athletes"

Detailed sport expertise scores of basketball players (Experts, n = 16)

. Weekly practice frequency was rated on a 4-point scale 1 = “from time to time”, 2 = “once per week”, 3 = “several times per week”, and 4 = “every day”. Competitive level was evaluated on a 4-point scale: 1 = Departmental level, 2 = Regional level, 3 = National level, and 4 = Feminine/Professional league.

| Participant | Weekly practice frequency | Years of basketball experience | Competitive level |
| --- | --- | --- | --- |
| 1 | 4 | 5 | 2 |
| 2 | 4 | 8 | 3 |
| 3 | 4 | 10 | 3 |
| 4 | 4 | 12 | 2 |
| 5 | 3 | 19 | 2 |
| 6 | 2 | 10 | 1 |
| 7 | 4 | 16 | 4 |
| 8 | 4 | 16 | 3 |
| 9 | 4 | 8 | 2 |
| 10 | 3 | 10 | 1 |
| 11 | 3 | 15 | 1 |
| 12 | 3 | 16 | 2 |
| 13 | 3 | 8 | 2 |
| 14 | 3 | 11 | 2 |
| 15 | 3 | 9 | 2 |
| 16 | 1 | 12 | 2 |
